# Effect of the Crystal Size of Biogenic Hydroxyapatites on IR and Raman Spectra

**DOI:** 10.1101/465146

**Authors:** S.M. Londoño-Restrepo, L. F. Zubieta-Otero, R. Jeronimo-Cruz, M. A. Mondragon, M. E. Rodriguez-García

## Abstract

This work focuses on the analysis of the impact that raw and calcined biogenic hydroxyapatite crystal size has on the Raman and infrared spectra. To this end, bovine, porcine, and human bones samples were defatted and deproteinized as well as calcinated at 720°C and then analyzed through Raman and Infrared spectroscopies, Transmission Electron Microscopy (TEM), Inductively Coupled Plasma (ICP), and Scanning Electron Microscopy (SEM). Raman and IR spectra for raw samples showed broad bands while after calcination bands became narrow and well defined. TEM images showed that all raw crystallites are nano-plates with a high crystalline quality contrary to the so far well-established concept that biogenic hydroxyapatites have low crystalline quality. This fact confirmed that the broad Raman and infrared bands of raw clean bones come from nanocrystal-plates. SEM analysis confirmed the increase in the size of the crystals after calcination from nano to sub-micron dimensions due to a coalescence phenomenon.

## INTRODUCTION

Nowadays bio-ceramic materials like hydroxyapatites are growing in importance in different fields, particularly in tissue engineering for medical and dental applications. The hydroxyapatite obtained from natural sources is called bio-hydroxyapatite (BIO-HAp), and its main difference with synthetic apatites is that it is a carbonated hydroxyapatite which contains other elements such as Na, Mg, Mn, Fe, among others (1, 2).

The physicochemical characterization of raw bone as well as BIO-HAp obtained through different methodologies is still an open problem due to the complexity of the material. Infrared and Raman spectroscopies have been extensively used to study these materials to monitor the removal of the organic matrix from the mineral phase as well as to identify different mineral phases and to study the changes in the crystalline quality of HAp caused by thermal processes (3, 4). However, no studies about the influence of the crystal size on the vibrational properties of synthetic and natural hydroxyapatites have been reported in detail. Usually, when these spectroscopies are used to study the vibrational states in synthetic and natural hydroxyapatites (5), the full width at the half maximum (FWHM) of a characteristic peak is used to determine the crystalline quality of HAp crystals and for clinical diagnostic (6). However, this criterion must be discussed in detail in the case of nanostructures.

It is well established in the literature that nanosized crystals produce wider Raman bands than micro-sized ones as can be found for semiconductors as Si (7, 8). The phonon confinement model has been proposed to explain Raman spectra in nanosized systems because the surface states must be considered (9-12). According to Gao et al. (13), the underlying mechanism behind the size-dependent Raman shifts is still quite controversial and an open problem. They proposed a theoretical method to explain the quantum confinement effects on the Raman spectra of semiconductor nanocrystals indicating that the shift of Raman bands in nanocrystals results from two overlapping effects: the quantum effect shift and a surface effect shift. Their calculations showed that there is a small shift in peak position as a function of the crystal size. On the other hand, they proposed this model for using Raman spectroscopy as a tool to measure size in nanostructures. Different works related to the Raman characterization of nano-synthetic HAp have been published in which the width of the band was not considered in the Raman interpretation. Concerning the analysis of the FWHM of Raman bands, Wopenka and Pasteris (14) showed that the significantly wider bands for biomaterials are indicative of the shorter-range disorder these nanocrystalline and carbonated phases have in comparison to those of synthetic hydroxyapatites. Even though they found that the studied biomaterials are in the nanoscale range, they did not consider that in nanoscale materials Raman dispersion is governed by surface states. The translational symmetry of the crystal is broken at grain boundaries which results in the appearance of specific surface and interface vibrational contributions (15). However, in the case of bio-hydroxyapatites, this information is still under discussion. For the determination of crystallinity in bones using Raman, the most used parameter is the width of the phosphate band at 959 cm^-1^ because it is an intense band with no overlapping from other bands (5, 16, 17). The calculation of the FWHM is carried out using a single Gaussian curve to fit the mentioned band. The fitted FWHM value is reported in wavenumbers, and its inverse is considered as a parameter that determines the crystalline quality of the samples. Nevertheless, this calculation does not make any physical sense when the particles are at the nano-scale.

FT-IR and Raman spectroscopy have also been used to quantify bone mineral crystallinity. Querido et al. (18) developed a methodology based on the ratio between selected bands to determine bone crystallinity. The X-ray diffraction peaks of the studied bones according to their interpretation had poor crystallinity thus resulting in broad and overlapping peaks. Nonetheless, this is misleading because it is well known that for nanoparticles, their X-ray diffraction patterns are formed by simultaneous and non-separable scattering and diffraction phenomena. Concerning the Raman analysis of the same samples, they found that the Raman bands varied in width and position due to the different degrees of crystallinity, but they did not perform the study of the crystalline quality of the samples by TEM.

The analyses of calcium phosphates by vibrational spectroscopies have been performed both from a theoretical and experimental point of view (19). According to De Mul et al. (17), the line broadening of the IR spectrum is related to the irregularity of the atomic array, referred to as lattice strain, among others. On the other hand, they consider that the crystallinity of the apatite domains can be determined by XRD. This kind of determination at a microscopic scale is difficult, so they chose a crystallinity index obtained from the broadening of the *v*_4_ apatite band at 600 cm^-1^. Here, it is essential to point out that the crystal size effects (finite crystal and surface states) are not considered. The same group made the calculation of the IR broadening with a relatively simple inter- and intra-ionic potential (Coulombian) in which all atoms have the same number of bonds, but the surface states break this rule and must be incorporated into the calculation.

Considering the above references, the aim of this work was to study the influence of the crystal size on the Raman and infrared spectra of hydroxyapatites from bovine, porcine and human bones to establish criteria to determine the size and crystalline quality of the synthetic HAp and BIO-HAp through FWHM measurements.

## MATERIAL AND METHODS

### Raw bone and annealed samples

Cortical bones from bovine, porcine, and human femurs were used for this work. Human femurs without any apparent pathologies were provided by the Universidad Autónoma de Querétaro while bovine bones (3-years-old) and porcine bones (157 days) were collected from the local slaughterhouse (folio number SDA-537295-98, 2017). All bones were defatted and deproteinized using the procedure proposed by Londoño-Restrepo et al., (1) in brief: the soft tissues of pre-cut 3 cm long bone slices were removed using a hydrothermal treatment with distilled water in an All Americans 1915X autoclave at 127 °C and 1.5 atm for 40 min three times. Subsequently, bones were dried off at 90 °C for 48 h to facilitate the grinding process. These bones were pulverized and sieved in a US 200 mesh (75 μm) to obtain powders with particles smaller than 75 μm. Finally, the bone powders were subjected to an alkaline hydrothermal treatment in a Ca(OH)_2_ solution (Mallincrodt Baker CAS No 1305-62-1) to remove the protein. Raw samples were labeled as B-Raw for bovine, P-Raw for porcine, and H-Raw for human. Afterward, the samples were incinerated at the same time using a Felisa (Mexico) furnace at 720 °C at a heating rate of 6 °C/min and a sintering time of 10 min as is shown in the thermal profile (Fig. 1) and later cooled inertially into the furnace in presence of air (20, 21). The annealed samples were labeled as B-720 for bovine, P-720 for porcine, and H-720 for human. On the other hand, a synthetic sample of hydroxyapatite from Sigma Aldrich was used in this study for comparative propose.

**Figure 1.**
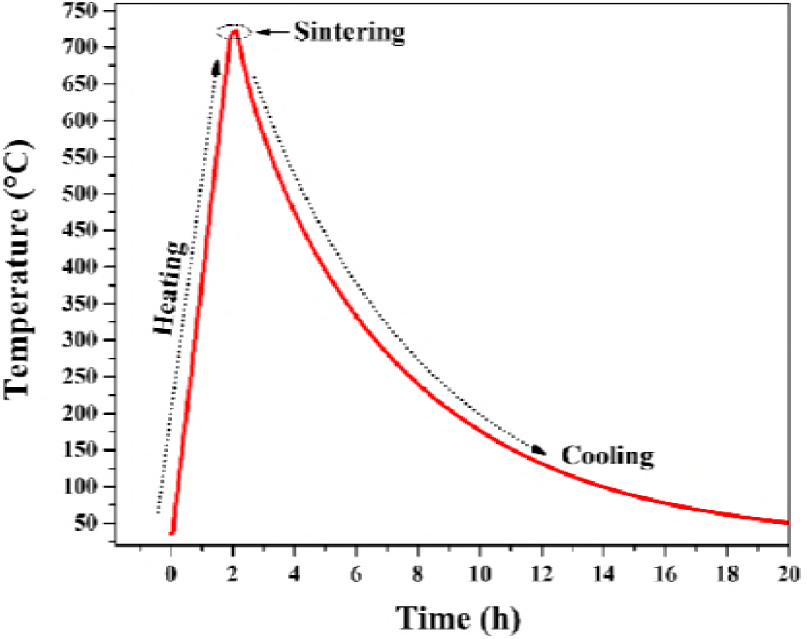
Thermal profile for powder bones incinerated at 720°C (B-720, P-720, and H-720).

### Mineral content: ICP

To determine the dependence of the vibrational spectra of raw bones on the different chemical elements they may have, the mineral composition of the untreated and calcinated samples was determined. Thus, the elemental composition of the BIO-HAp from human, bovine, and porcine bones, was determined with the use of a Thermo Fischer Scientific ICAP 6000 Series equipment with an argon plasma. 0.1 g of each sample was digested with 7 mL of nitric acid (Baker 69.3 %) using a temperature program; samples were analyzed in duplicate. Afterward, the samples were filtered (Whatman No. 42) and their volume was completed to 100 mL with deionized water. Finally, the samples were exposed to the argon plasma to excite the elements in the samples to identify them through their characteristic emission spectra. These were converted to elemental content by comparison to standard curves.

### Infrared spectroscopy

Infrared spectroscopy was used to follow the cleaning process aimed to defat and deproteinize the bones until the complete removal of the organic phase as well as to determine the influence of the crystal size on the infrared spectra of raw samples and samples incinerated at 720°C. The measurements were performed on a 6700 FTIR Thermo Scientific spectrometer equipped with an ATR (Attenuated Total Reflectance) accessory with a diamond crystal in the spectral range of 400 to 4000 cm^-1^ at a spectral resolution of 4 cm^-1^.

### Raman spectroscopy

All bone samples were analyzed using a Senterra Raman spectrometer from Bruker, equipped with a 785 nm laser and an Olympus microscope. A 20X objective was used, the spectral range measured was from 70 to 3500 cm^-1^, with a resolution of 3 cm^-1^ and the following instrument parameters: a 50 μ aperture, 100 mW of laser power of, 6 s integration time and 6 scans.

### TEM characterization

TEM images of B-Raw, P-Raw, and H-Raw were obtained. A high-resolution transmission electron microscope (S) TEM (JEOL ARM200F) was used to determine the crystal size of the bio hydroxyapatite samples; the acceleration voltage was 200 kV.

### Morphological studies of incinerated samples: SEM

Morphologic analysis of all BIO-HAp samples calcined at 720°C was carried out in a Jeol JSM 6060LV Scanning Electron Microscope, with an electron acceleration voltage of 20 kV. The samples were fixed on a copper holder with carbon tape and a gold thin film was deposited on them.

## RESULTS

### TEM characterization

Fig. 2 shows TEM images for defatted and deproteinized samples: P-Raw (A-D), B-Raw (E-H), and H-Raw (I-L). Here, it is very important to point out that according to Fig. 2 C, G, and K the bone from porcine, bovine, and human are formed by nano-plates. On the other hand, Fig. 2. B, F, and J clearly show that these BIO-HAps are in fact highly crystalline structures.

**Figure 2.**
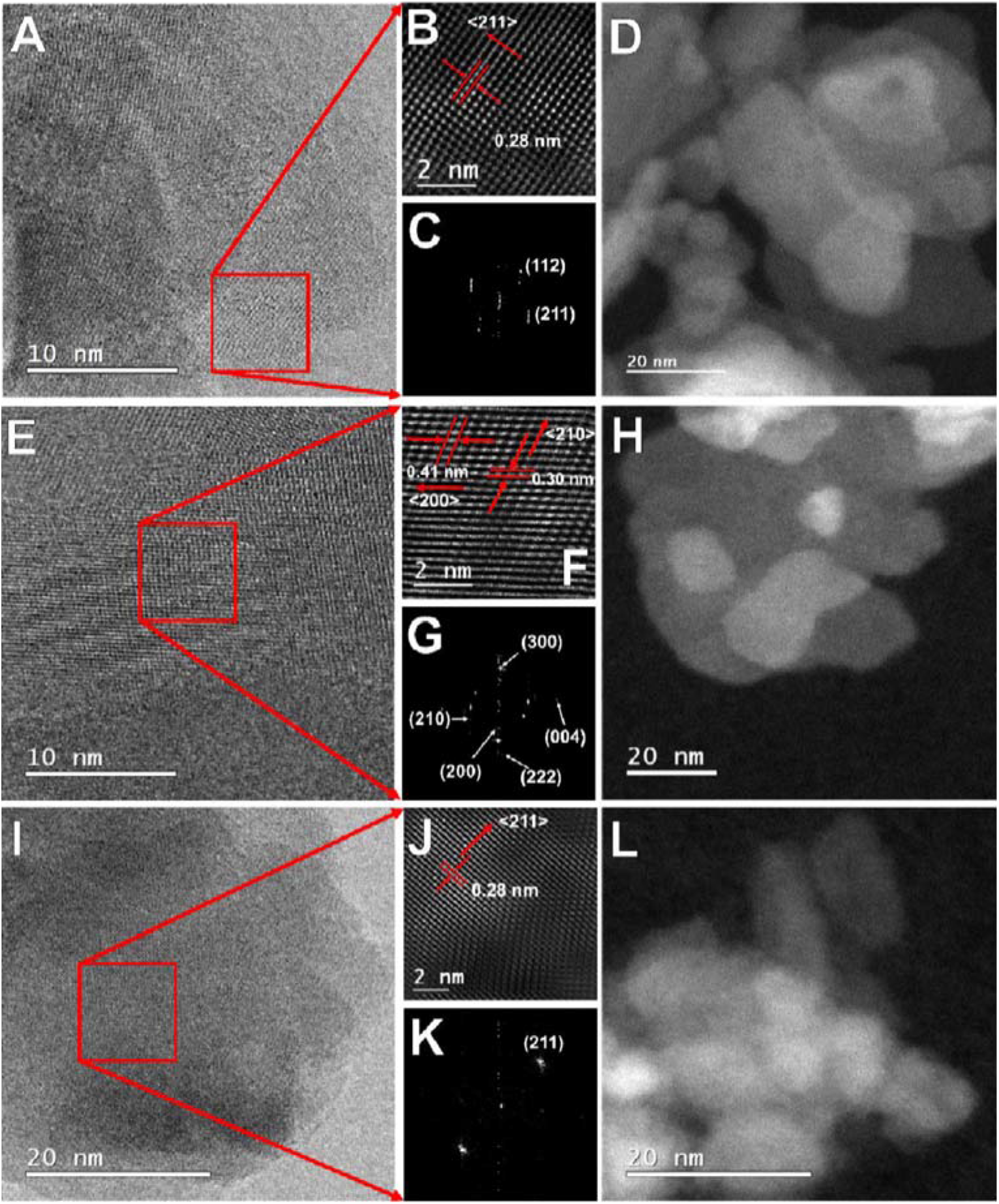
TEM images of the defatted and deproteinized porcine (A-D), bovine (E-H) and human (I-L) bones.

The images in Fig. 2 C, G, and K show that bones are formed by a polycrystalline arrangement of crystalline HAp nano-plates. The crystal size (2D) of these samples was determined using Image J software and are reported as the average value of 25 determinations. The crystal sizes obtained were: 21.35 ± 7.76 nm long and 6.03 ± 1.42 nm wide for human samples, 12.92 ± 2.55 nm long and 7.09± 1.05 nm wide for bovine samples and 16.53 ± 3.56 nm long and 6.37 ± 0.32 nm wide for porcine samples.

### Mineral composition by ICP

Fig. 3 A shows the P and Ca content and the Ca/P ratio for all samples: P-Raw, B-Raw, H-Raw, P-720, B-720, H-720, and synthetic HAp from Sigma-Aldrich. Ca percentage is from 21 to 37% while the P content is from 9 to 19%. Stoichiometric hydroxyapatite has 39.89% Ca and 18.5% P. In the same way, Ca/P ratio varies from 1.2 to 2.4 when is 1.67 for stoichiometric hydroxyapatite. Moreover, biogenic hydroxyapatites exhibit trace elements as Na, Mg, K, Fe, Al, and Zn with a maximum value of 9500 ppm (0.95%).

**Figure 3.**
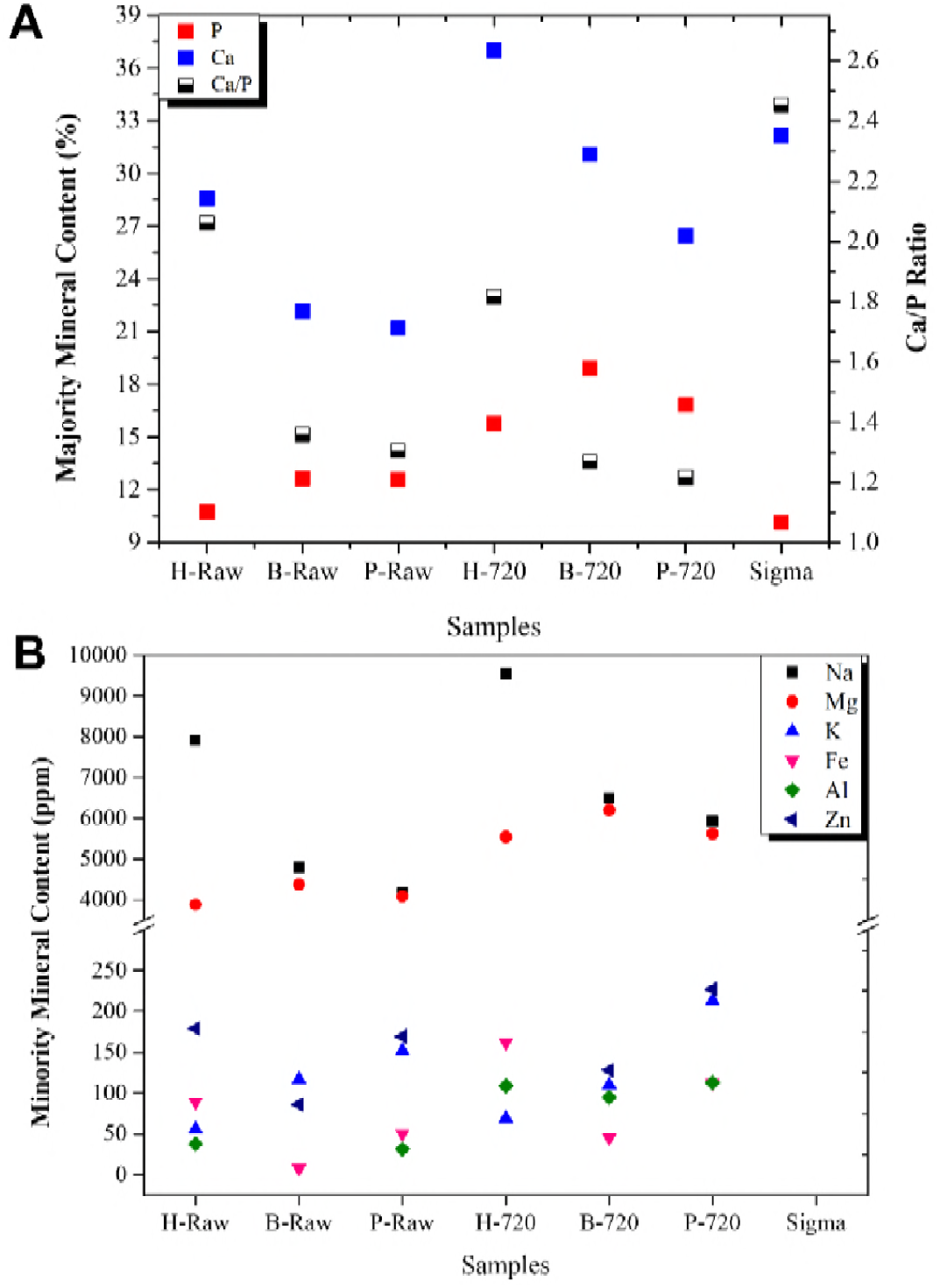
Mineral composition of defatted and deproteinized human, bovine and porcine powder bone. (A) Content of major minerals, P and Ca and Ca/P ratio. (B) Content of minor minerals: Na, Mg, K, Fe, Al, and Zn.

### IR vibrational analysis

Fig. 4 A to D shows the IR bands located between 400 to 4000 cm^-1^ for raw and calcinated samples, and Sigma Aldrich. The letters correspond to the identification carried out in this work, and these values were compared with others found in the literature in Table 1.

**Figure 4.**
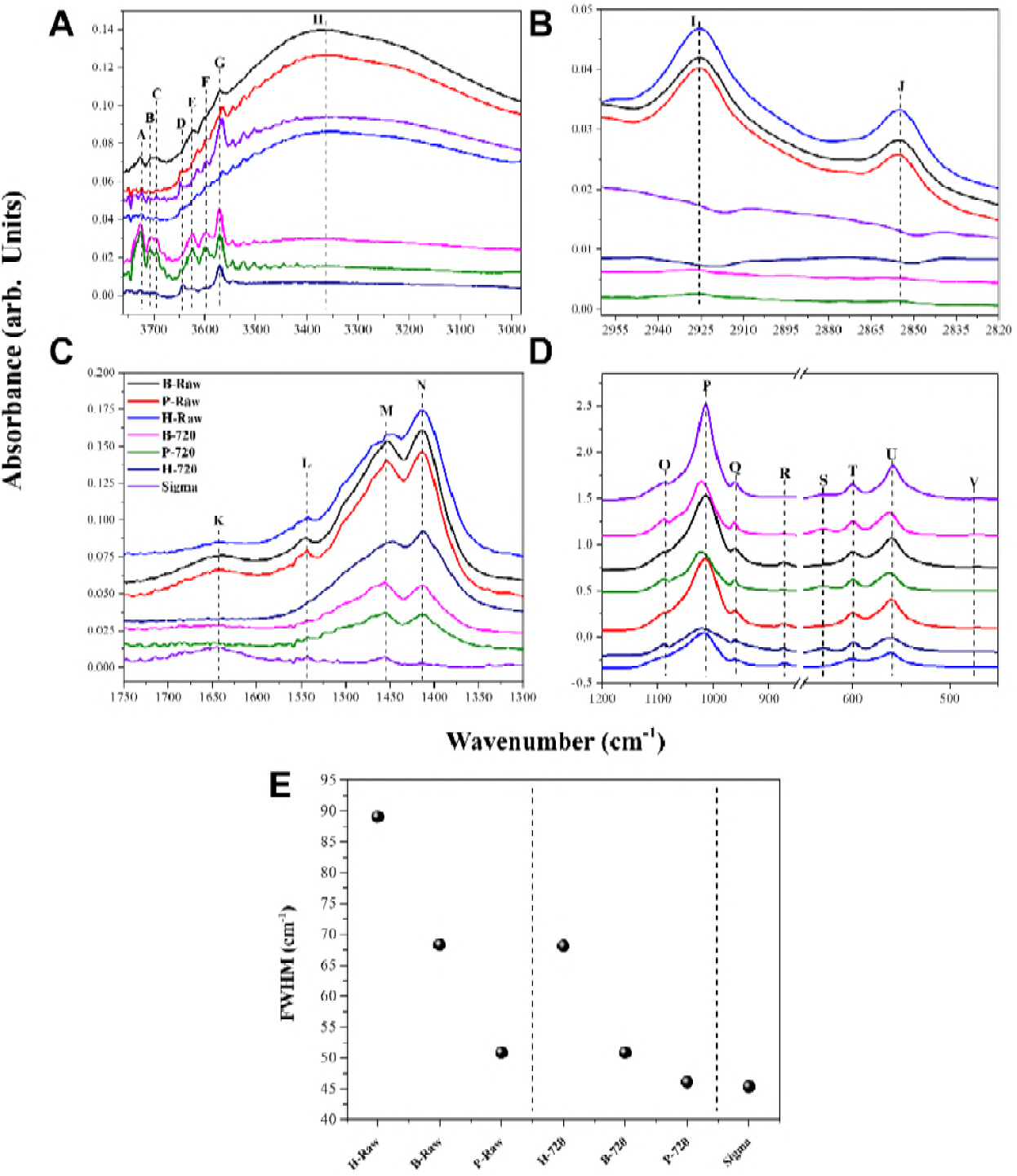
Infrared spectra of raw and calcined samples from human, bovine and porcine bones, as well as synthetic Hap, in the spectral ranges: (A) 3700-3000 cm^-1^, (B) 2955– 2820 cm^-1^, (C) 1750-1300 cm^-1^, (D) 1200-450 cm^-1^, and (E) FWHM of 560 cm^-1^ band for all samples analyzed.

**Table 1.**
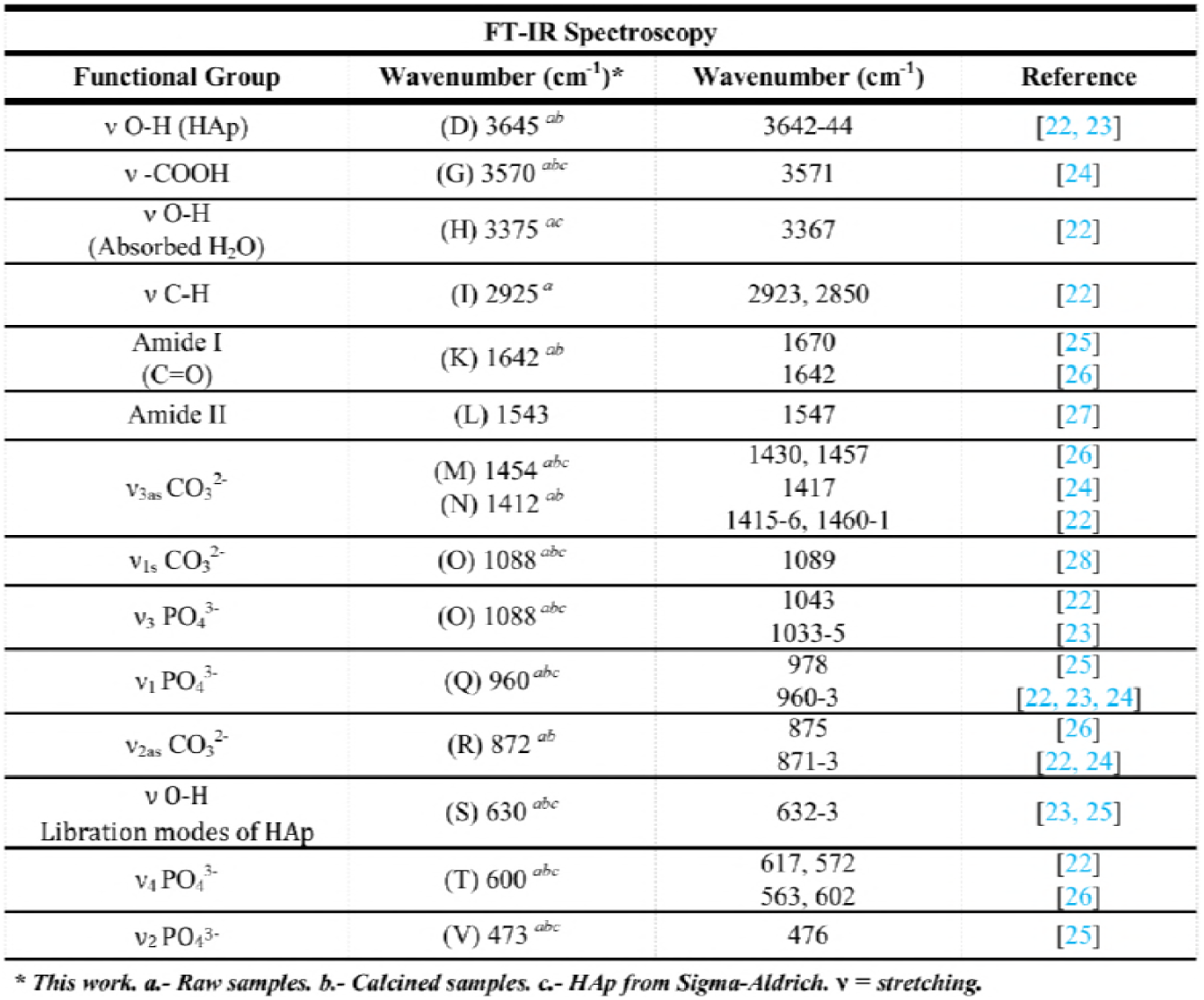
Observed infrared band positions for raw bones, sintered hydroxyapatites, and Sigma Aldrich.

Fig. 4. E shows the FWHM for raw and calcined samples, considering the band at 560 cm^-1^. After calcination, clearly the 560 cm^-1^ band becomes narrow and well defined. Some bands shown in Table 1 correspond to organic material and adhered water in the samples which disappear after calcination as H, I, J, K, and L bands (see Fig. 4 C). Regarding Sigma Aldrich sample, it does not exhibit any carbonate band then it is clear that this sample is a non-carbonated hydroxyapatite that contains a carbonyl group. The A, B, and C bands are presented in the studied samples. However, the origin of these bands is not understood.

### Raman vibrational analysis

The Raman spectra of the two sets of samples, raw and calcined, which correspond to nano- and micro-crystals, are displayed in Fig. 5, as for Sigma Aldrich hydroxyapatite. All found Raman bands are shown in Table 2.

**Figure 5.**
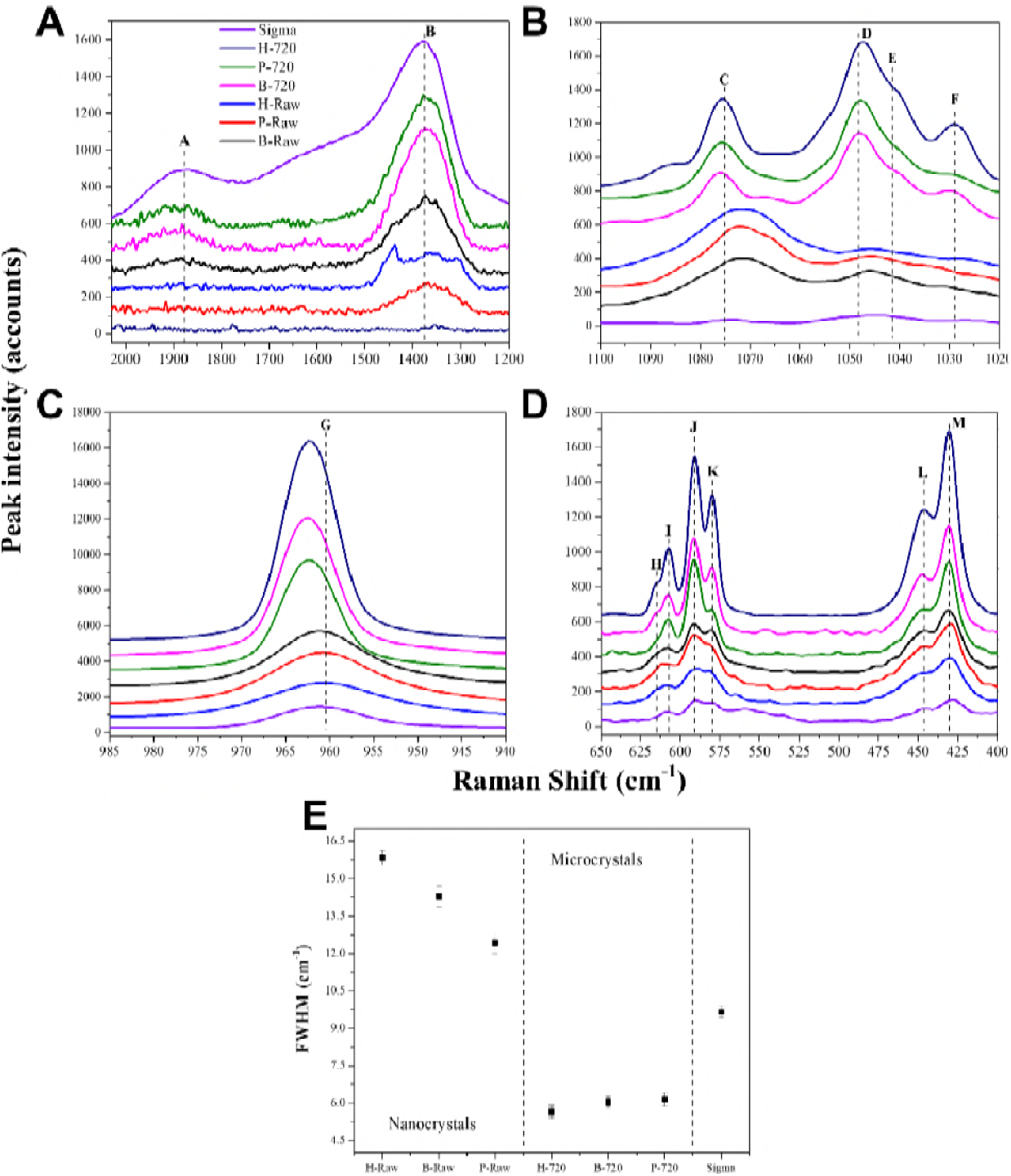
Raman spectra of raw and calcined samples from human, bovine and porcine bones, as well as synthetic Hap, in the spectral ranges: (A) 2000-1200 cm^-1^, (B) 1100-1020 cm^-1^, (C) 985-940 cm^-1^, (D) 650-400 cm^-1^, and (E) FWHM of 960 cm^-1^ band for all samples analyzed.

**Table 2.** Observed infrared band positions for raw bones, sintered biogenic hydroxyapatites, and Sigma Aldrich.

| RAMAN Spectroscopy |  |  |  |
| --- | --- | --- | --- |
| Functional Group | Wavenumber (cm<sup>-1</sup>)* | Wavenumber (cm<sup>-1</sup>) | Reference |
| Proteoglycan | (B) 1377 <sup>abc</sup> | 1375 | [29] |
| ν<sub>3</sub> PO<sub>4</sub><sup>3-</sup> | (C) 1076 <sup>abc</sup> | 1076 | [16, 30] |
|  | (D) 1048 <sup>b</sup> | 1045-8 | [20, 30, 31, 32] |
|  | (E) 1041 <sup>b</sup> | 1035, 1041-3 | [29, 31] |
|  | (F) 1028 <sup>b</sup> | 1030 | [20, 31] |
| ν<sub>1</sub> PO<sub>4</sub><sup>3-</sup> | (G) 960 <sup>abc</sup> | 960-1<br>955-7 | [20, 30, 31, 33] |
| ν<sub>4</sub> PO<sub>4</sub><sup>3-</sup> | (H) 614 <sup>ab</sup> | 579, 591 | [20, 30] |
|  | (I) 607 <sup>abc</sup> | 581 | [31] |
|  | (J) 591 <sup>abc</sup> | 587, 609 | [20, 32] |
| ν<sub>2</sub> PO<sub>4</sub><sup>3-</sup> | (L) 446 <sup>ab</sup> | 430-1, 447-50 | [30, 32] |
|  | (M) 430 <sup>ab</sup> | 425 | [20, 31] |

\*This work. a.- Raw samples. b.- Calcined samples. c.- HAp from Sigma-Aldrich. ν = stretching.

As can be observed from this figure, raw samples for porcine, bovine, and human bones have lower intensities and wider bandwidths as compared to those for the corresponding calcined bone samples; the FWHM value for the synthetic HAp sample lies between the values for the micro and nano samples.

### SEM analysis

The effect of the incineration process on the crystal size of the calcined samples is showed in Fig. 6 for P-720 (A), B-720 (B), and H-720 (C) samples. As was mentioned above, these samples were calcinated at 720°C using the same thermal profile. Fig. 2 showed that the raw samples are nanoplates with dimension around 20 nm length and 5 to 7 nm thick. After calcination, these samples get the following sizes: 70.38 ± 24.06 nm length and 45.52 ± 9.82 nm width for porcine sample, 52.31 ± 15.16 nm length and 36.58 ± 7.76 nm width for bovine sample, and finally 188.89 ± 74.53 nm length and 173.33 ± 48.25 nm width for human.

**Figure 6.**
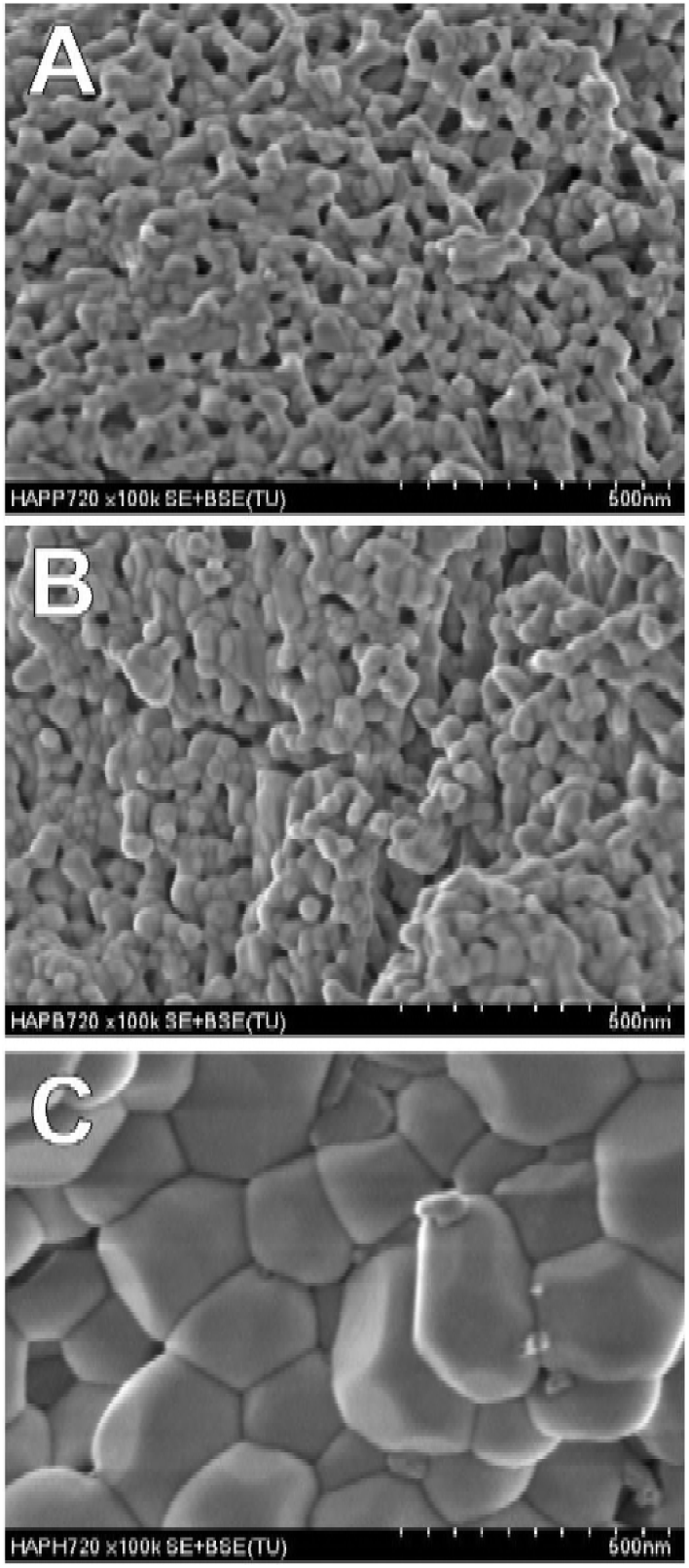
SEM image s of (A) P-720, (B) B-720, and (C) H-720.

## DISCUSSION

Results obtained from TEM analysis confirmed the nano-dimensions of the hydroxyapatite crystals of raw bones, which are forming longitudinal nano-plates, which can be correlated with the well-known longitudinal and preferential growth of the bone (1). TEM analyses showed the presence of well-defined nanocrystals and the (112) and (002) planes for bovine bone, (211) for porcine, and (112) for human, thus indicating a high crystalline order for all bone samples. Patel et al. (34) pointed out that raw bone powder from bovine is crystalline based on TEM images. As was mentioned before, the nano-sized dimensions of the hydroxyapatite crystals from all these bones, have a strong influence on the vibrational properties of bone.

Biogenic hydroxyapatites are no stoichiometric as was demonstrated through ICP, but this is not a disadvantage since these traces help to osteoinductivity when this material is used for medical applications. Ca levels for human samples are higher than for bovine and porcine due to the age of the bones where the samples were obtained; bovine and human samples correspond to adults while porcine is from young animals (5 months old). Other factors as diet, gender, and exercise can also modified their chemical composition (35). The difference in Ca and P levels between raw bones and calcined samples is due to a concentration effect since raw bone also contain small quantities of organic material. This fact directly affects the Ca/P ratio that is 1.67 for stoichiometric hydroxyapatite, as can be seen in Fig. 3 A. In the case of Sigma-Aldrich hydroxyapatite sample, it was found that its calcium value is the highest and its Ca/P ratio is higher than for a stoichiometric hydroxyapatite. Fig. 3 B exhibits the content of the minority elements in the bone which can be located as substitutional or interstitial atoms. Physicochemical properties, in special the structural and vibrational ones of biogenic HAp are affected by the presence of these ions because they modify the chemical environment which in turn modifies the dipolar moment and the polarizability. Here, it is worth noting that Sigma-Aldrich sample does not contain other minority elements.

As was showed before in Fig. 2, hydroxyapatite crystals from porcine, bovine, and human bones are constituted as nano-plates with sizes that vary from 5 to 30 nm length and from 5 to 7 nm width. The calculations to obtain the vibrational frequencies of a crystalline system assumes an infinite crystal size. However, in the case of nano-crystals, some atoms are located within the crystal and their coordination number on average is the same, while atoms on the surface exhibit different coordination numbers. Considering this fact, it is expected that the characteristic bands for the vibrational transitions producing the Raman and infrared spectra of hydroxyapatites suffer changes in position and width as compared with crystals.

Due to the annealing process, the nanocrystals of porcine and bovine bones suffer a coalescence process (20, 35) increasing their size. In this case, the new particles are from 52 to 70 nm in length and 36 to 45 nm in width. The calcination process for the human samples does not follow the same trend, since for these, the nanocrystals coalesce forming sub-micron crystals 190 nm length and 173 nm width. The difference in size the crystals reached after the annealing process for bovine and porcine samples as compared to that obtained for human bones is related to their respective chemical composition and age. Rendon and Serna (36) studied the effect of the crystal size on the infrared spectra of hematite formed between 250 to 600 °C; they showed that the differences in the infrared spectra are originated by variations in the size and shape of the particles from nanoparticle to microparticle systems. Through a detailed analysis of the IR bands, the bandwidth of bovine, porcine, and human raw samples exhibited different behaviors in comparison to calcinated samples. This effect is originated in part by the growth of the crystals size and to the calcination process changing the structural properties of these BIO-HAp (20, 35). Here is important to point out that if the crystal size increases, the number of vibrational states on the surface decreases, then the FWHM of the samples is not influence by these states and becomes narrow and well defined. Furthermore, taking into consideration the arguments stated above on the difference in coordination number for atoms in the crystals and on the surface, the position and width of Raman bands are expected to change when crystals of nano size and micro size are analyzed and compared.

The Raman FWMH measured showed a clear difference between nano and micro-sized crystals. The model to calculate the Raman spectra of a material assumes that the crystal is infinite, however advances in nanoscience and nanotechnology have made necessary to include finite size effects on the Raman and IR spectra to explain their features. The interpretation of the spectra plays a primary role in the case of nanoparticles as quantum dots (7, 8). The analysis of the experimental Raman spectra is done considering the confinement effect due to the finite size of the nanoparticles. The broadening of phonon peaks is governed by factors such as varied sizes, orientation, and shapes of the nanoparticles. Traditionally high FWHM values have been associated to crystals with low crystalline quality. However, in the case of biogenic hydroxyapatite nanocrystals, the high values they show are associated to size effects. Here, it is necessary to point out that the decrease in the *v*_1_ PO_4_^3-^ FWHM due to the incineration process is due to an increase in the crystallites size as was demonstrated by Londoño-Restrepo et al. (1) and measured in this work through SEM, as is shown above. Thus, it is not the result of the improvement of the crystalline quality of the HAp crystals, as it is frequently mentioned in the literature.

## CONCLUSIONS

Infrared and Raman spectroscopy were used to analyze the crystalline structure of raw and calcined at 720 °C of human, bovine and porcine bones. We have shown that defatted and deproteinized raw human, bovine, and porcine bones exhibit crystalline hydroxyapatite structures as nanoplates displaying a high crystalline quality, as opposed to common claims about the low crystalline quality biogenic hydroxyapatites have. This fact was made evident by TEM measurements. Besides, a calcination process at 720 °C to human, bovine and porcine bones gave, as a result, the formation of sub-micron crystals, and the infrared and Raman spectra of raw and calcined samples displayed marked differences. Considering that raw samples are nanoplates with high crystalline quality, the calculation of the crystalline percent in the present form does not make any physical sense.

## Author Contributions

All authors have contributed significantly to the manuscript. Study design: S.M.L.R. and M.E.R.G. Acquisition and analysis of data: M.A.M.S., L.F.Z., and R.J.C. Interpretation of data: All authors. Drafting of the manuscript: M.E.R.G. and S.M.L.R. Critical revision of the manuscript for important intellectual content: All authors.

